# Genome-wide association analysis of resistance to scald in an adapted multiparent winter malting barley population

**DOI:** 10.64898/2026.03.12.711358

**Authors:** J.M. Kolkman, S.S. Sepp, K. Kunze, G.C. Bergstrom, M.E. Sorrells

**Affiliations:** Plant Breeding and Genetics Section, School of Integrative Plant Science, Cornell University, Ithaca, NY, USA; Plant Pathology and Plant-Microbe Biology Section, School of Integrative Plant Science, Cornell University, Ithaca, NY, USA

## Abstract

Scald, caused by the fungus *Rhynchosporium graminicola* Heinsen 1897, is a major foliar disease in winter malting barley (*Hordeum vulgare* L). Resistance to scald in winter malting barley is controlled by major and minor resistance genes. We used a population of 377 lines derived from biparental crosses among five winter malting barley parents to analyze resistance to scald and associated agronomic traits. Increased winter survival and later heading dates were negatively correlated with increased resistance, whereas increased height was positively correlated with resistance. A genome-wide association study (GWAS) for resistance to scald was analyzed with multiple models, using 14,789 SNP and 374 lines. The similarities and differences between the models were identified in SNP trait associations and phenotypic effect sizes. SNP associations identified a large region on chromosome 3H across models. FarmCPU identified additional associations on chromosomes 2H, 3H, 4H and 7H. Linkage disequilibrium on chromosome 3H and GWAS for resistance to scald using the *Rrs1*-linked marker, HVS3, as a covariate confirmed *Rrs1* was segregating in this population. GWAS for winter survival, heading date and plant height identified associations across the genome, with chromosome 2H showing SNP-trait colocalizations between resistance to scald, winter survival, heading date and plant height. Breeding for durable resistance to scald in winter malting barley can include pyramiding major resistance loci, such as *Rrs1*, as well as QTL for disease resistance and agronomic traits.

**PLAIN LANGUAGE SUMMARY:** *Genetic architecture of resistance to scald in winter malting barley:* Scald is an important foliar pathogen in winter malting barley, affecting both grain yield and quality. While resistance to scald is controlled by major and minor resistance genes, agronomic traits are also known to limit the spread of scald in barley. We determined the genetic architecture using a large multiparent population of winter malting barley. The FarmCPU genome-wide association model proved optimal for defining the resistance genes, with the major resistance gene, *Rrs1*, conferring 29% of the variation in this population. Fewer days to heading and taller plants contributed to plant avoidance of scald. Reduced canopy coverage in plants with low winter survival led to less scald severity. A region of the genome contributing a minor resistance effect was co-localized with a region for plant height, heading date and winter survival.

**Core Ideas:**

- Resistance to scald in a large multiparent population was derived from a major resistance gene (*Rrs1*) and several smaller effect QTLs
- *Rrs1* resistance was derived from ‘Lightning’ and is located within a large linkage block on Chromosome 3H
- Fewer days to heading and taller plants were correlated with less disease in the multiparent winter malting barley population in NY state
- A QTL for resistance to scald co-localized on chromosome 2H with winter survival, heading date, and plant height
- FarmCPU was an optimal model for association analysis for resistance to scald in the multiparent unbalanced diallel population.

## 1 INTRODUCTION

Barley, *Hordeum vulgare* L., is a globally important crop used for livestock feed, human food, and for malt used in brewed and distilled beverages. The increased demand for malting barley by the craft brewing industry has led to increased malting barley production in non-traditional barley growing regions (Shrestha & Lindsey, 2019). The adoption of malting barley in the Northeast US has been fostered by development of improved management recommendations for barley grown in higher rainfall and more humid conditions than in traditional barley growing regions (Shrestha & Lindsey, 2019; Siller et al., 2021). Winter barley offers consistently higher yield and often higher quality than spring barley and often avoids unfavorable spring planting conditions and summer droughts. Along with agronomic management differences, a shift from spring to winter malting barley alters the disease exposure profile, resulting in an environment conducive to the proliferation of scald. Barley scald is a foliar fungal pathogen, that flourishes in the cooler climates reaching up to 77% diseased leaf area of upper canopy during early grain filling stages in susceptible cultivars grown in New York (Kolkman et al., 2025a). Scald can cause both a reduction in quality (Avrova & Knogge, 2012), and yield losses ranging from 10 to 45% (Shipton et al., 1974), and up to 65% in severe epidemics (Beigi et al., 2013).

Scald in barley is caused by *Rhynchosporium graminicola* Heinsen 1897 (Crous et al., 2021; formerly known as *R. commune;* Zaffarano, et al., 2011) a hemibiotrophic foliar fungal pathogen that thrives in cooler temperatures (Zaffarano et al., 2008). Originally identified in 1897 (Frank, 1987), *R. graminicola* was renamed as *R. secalis* (Oudem.) J.J. Davis in 1919 (Davis, 1919), renamed again in 2008 as *R. commune* to indicate host speciation specific to *Hordeum* species (Zaffarano et al., 2008, 2011). *Rhynchosporium graminicola* is proposed to have originated in the cool climates of Scandinavia approximately 2500 years ago as a host jump from an unknown grass species (Brunner et al., 2007). *Rhynchosporium graminicola* spreads through previously infected barley debris and/or infected seed (Ababa et al., 2023). Conidia land on leaf surfaces and produce a germ tube along the intercellular grooves that produces an appressorium that penetrates through the cuticle, all within the first 24 hours of exposure (Linsell et al., 2011). Hyphae grow below the cuticle, above the anticlinal walls and between the epidermal cells, between the pectin layer and outer cell wall, disrupting the pectic layer and cuticle for pectin degradation (Ayesu-Offei & Clarke, 1970; Ryan & Grivell, 1974; Lehnackers & Knogge, 1990; Linsell et al., 2011). The cuticle and epidermal layers separate with the production of a packed hyphal mat, forming a subcuticular stroma, inducing a water-soaked lesion, all within approximately 4 to 8 days (Ayesu-Offei & Clarke, 1970; Linsell et al., 2011). Mesophyll collapse follows approximately 7 to 14 days post inoculation, creating a straw-colored necrotic lesion surrounded by dark brown borders at which point hyphal growth increased, likely due to the release of nutrients from the collapsed mesophyll cells (Ayesu-Offei & Clarke, 1970; Lehnackers & Knogge, 1990; Linsell et al., 2011). Conidia are produced from subcuticular and/or substomatal stroma, which protrude through the cuticle along the leaf surface.

Resistance to scald in barley has been attributed to both major resistance genes and quantitative trait loci (QTLs; as reviewed by Zhang et al., 2020). In total, 11 major resistance genes have been described (as compiled by Noe et al., 2025) with major resistance genes located on all chromosomes except chromosome 5H. Several of the *Rrs* genes are comprised of multiple alleles and/or loci, such as the *Rrs1* gene complex located on chromosome 3H that includes the previously described alleles *Rh, Rh1, Rh3 Rh4* and *Rh7* (Bjørnstad et al., 2002). While no causal gene underlying the *Rrs* genes has been determined, several *Rrs* genes have been fine-mapped. The *Rrs1* gene region has been delimited to a region of 0.8 Mbp with 10 candidate genes using the Morex genome sequence. The *Rrs1* gene is not, however, present in the Morex genome, and is hypothesized to be presence-absence and/or a gene duplication variant. There are several candidates that are plausible in this genomic region that is reticent to recombination, due to its centromeric location, including a protein kinase (Looseley et al., 2020). The *Rrs2* resistance gene is proposed to be in a cluster of pectin esterase inhibitor (PEI) genes, either as a unique gene, a combination of PEI genes and/or other unidentified gene(s) (Marzin et al., 2016). The *Rrs13* resistance gene, fine-mapped within a 0.58 to 1.2 Mbp region, codes for two to seven tandemly repeated leucine-rich repeat receptor-like proteins (LRR-RLP) and a lectin receptor-like kinase (Eckstein et al., 2024). More recently, the *Rrs18* gene has been identified upstream of *Rrs13* (Coulter et al., 2019). Fine-mapping and RNA expression analysis identified four candidate genes with the most likely candidate gene a serine/threonine protein kinase. As well as these four genes, a stable QTL for adult plant resistance to scald, QSc.VR4, was fine-mapped to a 0.38 Mbp region that included an LRR-RLK multi-gene family, and a germin-like protein multigene family (Wang et al., 2020).

Numerous genome-wide association studies (GWAS) have been utilized to identify and characterize resistance to scald in barley. Association studies have relied on a variety of populations, including diversity panels (Looseley et al., 2018; Hiddar et al., 2023; Kunze et al., 2024; Noe et al., 2025), a multiple advanced generation inbred cross (MAGIC) population (Hautsalo et al., 2021), the HEB25 nested association mapping population (Büttner et al., 2020), and a variety of populations that included diverse germplasm, including landraces, wild species and breeding lines (Looseley et al., 2018, 2020; Clare et al., 2023; Ababa et al., 2024; Ijaz et al., 2024). Many trials utilize greenhouse trials to ascertain seedling resistance (major gene resistance), using one or more isolates to characterize resistance to specific isolates. Several studies relied on (natural) field infection to identify adult plant resistance (Daba et al., 2019; Looseley et al., 2020; Ijaz et al., 2024). All GWAS studies used spring barley, except for two studies that used spring, winter and/or facultative barley lines (Looseley et al., 2020; Kunze et al., 2024). Adult plant resistance in the field encompasses natural infection, and resistance comprised of major resistance (*Rrs*) genes and/or QTL for resistance to scald. Adult plant resistance has been associated with genes coding for plant growth traits, such as *sdw1*, the gibberellin 2-oxidase gene, *HvGA20ox2* (Xu et al., 2017), implicated with plant height and susceptibility to scald (Looseley et al., 2018). Improving barley for resistance to scald relies on agronomic practices as well as breeding for resistance. Due to the potential for *R. graminicola* populations to overcome resistance (Mcdonald, 2015; Ababa et al., 2024), understanding the facets of resistance is imperative in breeding for sustainable resistance.

Genome wide association studies are useful for the identification of marker-trait associations and characterization of trait architecture across the genome. The premise of GWAS relies on maximizing meiotic recombination events by utilizing populations consisting of a large diverse number of genotypes in conjunction with high SNP marker density, and in many cases, using population structure and kinship to minimize false positive results (Flint-Garcia et al., 2005; Yu et al., 2006). Several models of GWAS have been developed that use different approaches to identify marker trait associations (Tibbs Cortes et al., 2021). In contrast, breeding programs are targeted to regional adaptation and are generally limited to elite material based on founders and the introgression of adapted material that may have originated from diverse or pre-breeding efforts (Kelly et al., 1998). Large populations are created with the goal of finding the transgressive segregants that move crop development and the release of novel cultivars forward.

In this study we characterized resistance to scald in an unbalanced diallel breeding population of winter malting barley recombinant inbred lines and doubled haploids derived from biparental crosses between five founder lines that are adapted to New York environments. We examined the correlation of agronomic traits, such as winter survival, days to heading and plant height with resistance to scald as a means of escape and/or avoidance mechanisms. Using GWAS, we characterized the genetic architecture of resistance to scald in the multiparent population, and the co-localization of SNP associations between resistance to scald and winter survival, heading date and plant height. We explore a suite of GWAS models available in GAPIT version 3 (Wang & Zhang, 2021) to determine the appropriate GWAS model for the multiparent unbalanced diallel population, given the relatedness and limited meiotic recombinant events between lines within the germplasm.

## 2 MATERIALS AND METHODS

### 2.1 Germplasm and trait phenotyping

#### 2.1.1 Plant germplasm

A population of 377 winter malting barley breeding lines was derived from an unbalanced diallel mating design using four winter malting barley cultivars, including ‘Flavia’,, ‘KWS Scala’, ‘SY Tepee’, and ‘WintMalt’ and one facultative malting barley cultivar (‘Lightning’; Hayes et al., 2021). F_1_ plants from the corresponding 10 cross combinations were advanced to the F_5_ stage through single seed descent and/or creation of doubled haploids resulting in 183 recombinant inbred lines (RILs) and 194 doubled haploid (DHs) lines (Figure 1) for a total of 377 lines (Figure 1). The largest number of lines was derived from ‘Lightning’ x ‘SY Tepee’ crosses, with 93 lines. ‘Lightning’ was the parent with the most lines represented, with 272 (72%) of the 277 lines. ‘WintMalt’ was the parent with fewest lines represented with only 95 of the 377 lines (25%). The fewest number of lines were from the cross between ‘WintMalt’ x ‘SY Tepee’, with four lines. Additional parents were present in the population with 43.8% (165 lines), 35% (132 lines), 26% (100 lines) for ‘SY Tepee’, ‘Flavia’, and ‘KWS Scala’ respectively.

**Figure 1.**
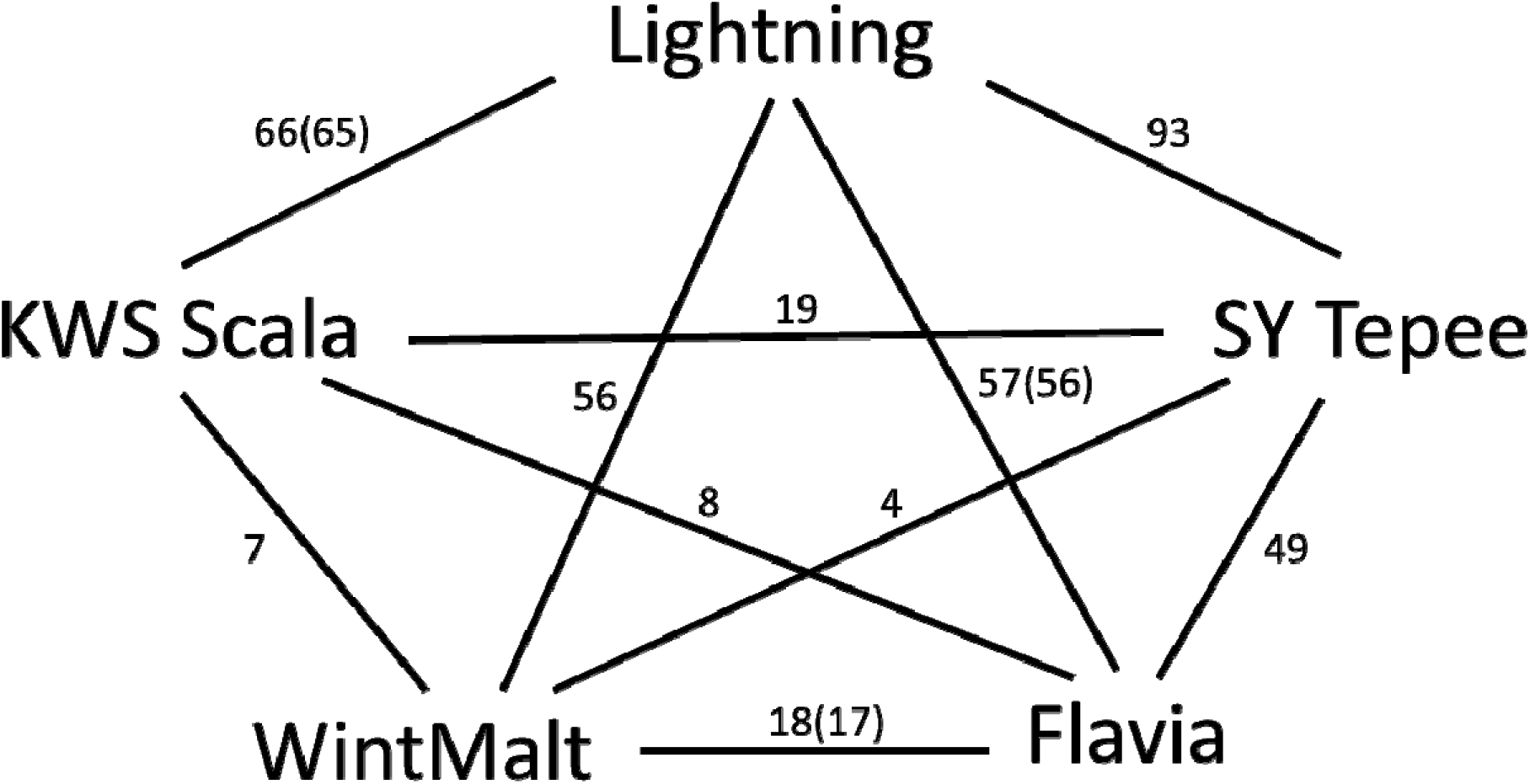
Structure of the unbalanced diallel multiparent population derived from five adapted cultivars. The 377 lines consisted of recombinant inbred lines and doubled haploids. The number of lines between each biparental cross is indicated along the line between the two parents, which were used for statistical analysis (377 lines) and for GWAS analysis (374 lines) with three lines less as indicated by the numbers in parenthesis.

#### 2.1.2 Field trials

The diallel population was planted in four field sites (environments) at the Cornell University Campus Area Farms in Tompkins County near Ithaca, New York and included the Snyder Farm and Helfer Farm field sites in 2022, and the McGowan Farm and Ketola Farm field sites in 2023. Trials were grown in an augmented design with a single replication in each environment and included the parental cultivars as well as check cultivars (‘KWS Scala’, ‘Lightning’ and ‘Endeavor’) within four blocks in both 2022 environments (Snyder Farm and Helfer Farm), and within eight and ten blocks in Ketola Farm and McGowan Farm field sites in 2023, respectively. Seeds were planted in plots with 1.2 m width, 3.0 m plot length, with a 17.8 cm space between rows within the plot and 25.4 cm between plots. No plant growth hormone or foliar fungicides were applied. Preplant fertilizer of 10:10:10 was applied at 224 kg/ha (22.4 kg ha^-1^ N) and followed by a top dress in the spring of 67 kg ha^-1^ N via liquid Urea-ammonium nitrate 30. The spring applied herbicide regime in 2022 consisted of Harmony (35 g ha^-1^), bromoxynil (BROX-2-EC at 1.5 L ha^-1^) and Induce surfactant (0.7 L ha^-1^). Herbicide applications in the spring of 2023 consisted of Axial XL (1.2 L ha^-1^), Harmony Extra SG (35 g ha^-1^) and Induce surfactant (0.7 L ha^-1^). No foliar fungicide was applied.

#### 2.1.3 Trait phenotyping

Scald infections relied on the natural inoculum present at each field site. *A priori* knowledge of these field sites indicated a historical presence of the scald pathogen in each environment. The scald susceptible cultivar ‘KWS Scala’ (Blachez et al., 2018) was planted in each block of each experiment and acted as a susceptible check and spreader ensuring adequate conidial inoculum across the trial. Plots were scored for scald symptoms at approximately Feekes stage 11.1 to 11.3 on June 13th and 9th in 2022 at the Helfer Farm and Snyder Farm field sites, respectively, and at Feekes stage 11.1-11.3 from June 26^th^ to the 30^th^, and June 15^th^ to 19^th^ in 2023 at the McGowan Farm and Ketola Farm field sites respectively. Plots were scored on a 0-9 scale in all four environments. Severity was also scored as a percentage diseased leaf area (DLA) for the upper canopy in 2023. 2022 trial scores were converted to a % DLA. Additional agronomic traits were scored in the trials including winter survival (WS), measured as percentage of plants surviving taken in the early spring; heading date (HD), when 50% of the heads had completely emerged from the sheath (converted to Julian heading date); and plant height, measured as the height (HT) from the ground to the top of the spike excluding awns.

#### 2.1.4 Data analysis

The scald DLA, WS, HD and HT were analyzed separately in 2024 JMP® 17.1 (JMP Statistical Discovery LLC, Cary NC) using a mixed model (restricted maximum likelihood; Reml) with genotype as fixed effects, and environment, row within environment, and column within environment as random model effects. Least squared means were used to determine frequency distributions and parental phenotypic value in comparison to the population. Spearman’s correlations between DLA, WS, HD, and HT were determined using JMP. Heritability was determined on an entry-means basis where *h*^2^ = σ^2^_g_/(σ^2^_e_/t+ σ^2^_g_), with σ^2^_g_, and σ^2^_e_ representing genetic variance, and experimental error, respectively, and t representing number of test environments (Fehr, 1987).

To normalize the residuals and homoscedasticity for association analysis, the DLA, WS, HD, HT measurements were analyzed in a simple linear model in R to estimate the residual parameters for the population of 377 without controls, with model effects of genotype, environment, row within environment and column within environment as fixed effects. A constant of 1 was added to each DLA measurement to avoid nulls for transformation. The best lambda for transformation of the data was determined using the Box-Cox function (Box & Cox, 1964) in the MASS package in R v3.2.3 (R Core Team, 2015), that was used to transform for normal residuals and homoscedasticity. The DLA (with added constant of 1), WS, HD and HT were transformed using respective lambda values of -0.1818, 2.9494, - 7.0303 and 0.0606 for DLA, WS, HD and HT, respectively. The HD transformed data was multiplied by 1 x 10^16^ for further analysis.

The best linear unbiased predictors (BLUPs) transformed DLA scores were estimated using Reml in JMP with WS, HD and HT as fixed covariate effects and genotype, environment, row within environment, and column within environment as random effects. The BLUPs for the transformed values for DLA, WS, HD, and HT were also calculated using Reml in JMP with genotype, environment, row within environment and column within environment as random effects.

### 2.2 Genotypic Analysis

#### 2.2.1 SNP Genotypic Data

Plant tissue was harvested from the multiparent population at the two-leaf stage. Tissue was lyophilized and a modified Cetyltrimethylammonium bromide (CTAB) extraction was used for DNA extraction (Doyle & Doyle,1987). DNA was genotyped with the 50K Illumina iSelect SNP array (Bayer et al., 2017; Mascher et al., 2017, 2021) at the USDA-ARS North Central Small Grains Genotyping Lab in Fargo, ND and resulted in high quality SNP data for 374 lines. The SNP data was trimmed from the original 43,078 SNPs to exclude homozygous SNPs across the population using TASSEL 5.0 (Bradbury et al., 2007). The 15,463 SNPs were used to estimate population structure.

#### 2.2.2 Marker Genotypic Data

Leaf tissue was collected for specific marker traits associations (MTA) within known resistance and/or height genes. For the MTA analysis, three kernels per line included in the multiparent population were planted in a 96 cell tray and grown under a light bench. Tissue was harvested at the seedling stage and lyophilized. The modified CTAB extraction was used to extract DNA (Doyle & Doyle, 1987).

The multi-parent population was genotyped for the plant height gene, *HvGA20ox2*, also known as *semi-dwarf1* (*sdw1*), to test for the presence of the *sdw1.d (Diamant)* or *sdw1.c* (denso) alleles (Xu et al., 2017). The *sdw1.d* and *sdw1.c* alleles were amplified via polymerase chain reaction using mutant specific primers based on Xu et al. (2017), with the adaptation of a fluorescent M13-tail in the forward primers for both the sdw1.d allele (5401F_M13F; 5’-TGTAAAACGACGGCCAGTGGTGCTCCAGACCGCTCAG-3’) and sdw1.c (MC40861P3F_M13F; 5’-TGTAAAACGACGGCCAGTTATGGCGTGACCAAAGGTTC-3’) that correspond to the reverse primers for *sdw1.d* (5549R; 5’-CGGCGGAGGGGTCAATG-3’), and *sdw1.c* (MC40861P4R; 5’-CACCAATCCACCACGAAGA-3’). The PCR reaction consisted of ∼30 ng genomic DNA, 12.5 µl of GoTaq polymerase, 0.8 µM M13F primer, 6 µM R primer, 6 µM M13-FAM primer (*sdw1.d*) or M13-VIC primer (*sdw1.c*) and 8.22 µl of H_2_0 in a 25 µl reaction. The PCR amplification protocol for the sdw1.d primer pair included an initial denaturation step of 94°C for 3 m, 30 cycles (94°C for 1 m, 55°C for 30 s, 72°C for 30 s), 10 cycles (94°C for 1 m, 50°C for 30 s, and 72°C for 30 s), and a final extension cycle of 72°C for 20 m. The PCR amplification cycle for the sdw1.c was similar to above, however the annealing temperatures in the two cycles were 54°C and 53°C. Fragment analysis was performed on pooled samples by the Biotechnology Resource Center (BRC) Genomics Facility (RRID:SCR_021727) at the Cornell Institute of Biotechnology *(http://www.biotech.cornell.edu/brc/genomics-facility*) on an Applied BioSystems 3730xl (Thermo Fisher Scientific, Waltham, MA), with the ABI 500LIZ size standard. Results were analyzed in Genemarker (SoftGenetics, LLC, State College, Pennsylvania).

In addition, the HVS3 SCAR marker (Genger et al., 2003) was screened in the parental lines and multiparent population using ∼20ng genomic DNA, 1X GoTaq ® Green Master Mix (Promega Cooperation, Madison, WI), 5µM Forward primer (5’-AAT CCT ACC TAT CCC ACC TT-3’), 5 µM Reverse primer (5’-TAT TTT CAG CCT TGT TCG GC-3’) in a final reaction volume of 25 µl. The DNA was amplified via PCR using an initial step of 94°C for 3 m, 35 amplification and extension cycles (94°C for 30 s, 50°C for 30 s, 72°C for 1 m), followed by a final extension cycle of 72°C for 20 m. The PCR products were amplified on a 2% agarose gel using GelRed ® Nucleic Acid Stain (Biotium, Inc, Freemont, CA) for verification of PCR on a UV lightbox.

### 2.3 Genome-wide association analysis

#### 2.3.1 Genome-wide association mapping for resistance to scald with multiple models

Association analysis was used to identify regions of the genome that conferred resistance to the scald pathogen in our adapted germplasm. Phenotypic data included the transformed DLA, WS, HD and HT BLUPs. Of the diallel population, 374 lines were used that included both genotypic and phenotypic data. The 43,078 SNPs (Morex version 3) derived from the 50K Illumina iSelect SNP array (Bayer et al., 2017; Mascher et al., 2017, 2021) were filtered to exclude homozygous SNPs across the population in TASSEL 5.0 (Bradbury et al., 2007).

Genome-wide association for Box-Cox transformed DLA BLUPs for scald that included WS, HD and HT as covariates as previously described was ascertained using GAPIT version 3 (Wang & Zhang, 2021) in R Studio (RStudio 2025.09.1+401, Posit Software, PBC). The Mixed Linear Model (MLM), Multiple Locus Mixed Linear Model (MLMM), Fixed and random model Circulating Probability Unification (Farm CPU) and (Bayesian-information and Linkage-disequilibrium Iteratively Nested Keyway (BLINK) models (Yu et al., 2006; Zhang et al., 2010; Segura et al., 2012; Liu et al., 2016; Huang et al., 2019), using the 14,789 SNPs, were used to estimate marker trait association parameters for resistance to scald. Kinship and principal component analysis (PCA) were determined within GAPIT version 3 to be used in the appropriate models. The Bonferroni threshold (Bonferroni, 1936) was used as an initial default significance threshold for *R*^*2*^ (MLM) and Percent Variation Explained (PVE; MLM, MLMM, FarmCPU, and BLINK). The resulting Quantile-Quantile (QQ) plots and contributing PVE were examined to determine the appropriate GWAS model for this structured population. To reduce Type II false negative errors due to the conservative nature of the Bonferroni threshold, the QQ plots were examined to determine the best GWAS fit that combined the reduction of spurious associations and putative significant or strong associations. The MLMM, FarmCPU and BLINK GWAS model QQ plots showed a *p*-value differentiation at the -log_10_(*p*) = 4.0. To determine the appropriate *R*^*2*^ (MLM only) and PVE (MLM, MLMM, FarmCPU and BLINK models) the ‘N.sig = n’ prompt was used in GAPIT version 3 for the significance threshold where ‘n’ equals the number of markers for each model equal or above the -log_10_(*p*) = 4.0 threshold. An additional association analysis using FarmCPU with PCA = 3 as a covariate was used to validate model selection with or without PCA.

#### 2.3.2 The *Rrs1* gene complex

Linkage disequilibrium (LD) surrounding the two strongest SNP associations identified on chromosome 3H at 174 Mbp and 442 Mbp were analyzed to determine if they were the same peak or independent associations. The LD was first assessed in TASSEL 5.0 (Bradbury et al., 2007) between the SNP at 174 Mbp and all the other SNPs on chromosome 3H (Figure 3A), and between the SNP at 442 Mbp and all the other SNPs on chromosome 3H. The Manhattan plots for the GWAS MLM model (Figure 2) were overlaid with LD estimates and used to confirm the presence of a large linkage block across the centromeric region of chromosome 3H, as visualized in the ‘ggplot2’ package of R v3.2.3 (R Core Team, 2015). The GWAS MLM model was used to assess associations using GAPIT version 3, using the same methodology as above, except for only using the 354 lines in the GWAS model and using the HVS3 marker data as a covariate (Figure 3C).

**Figure 2.**
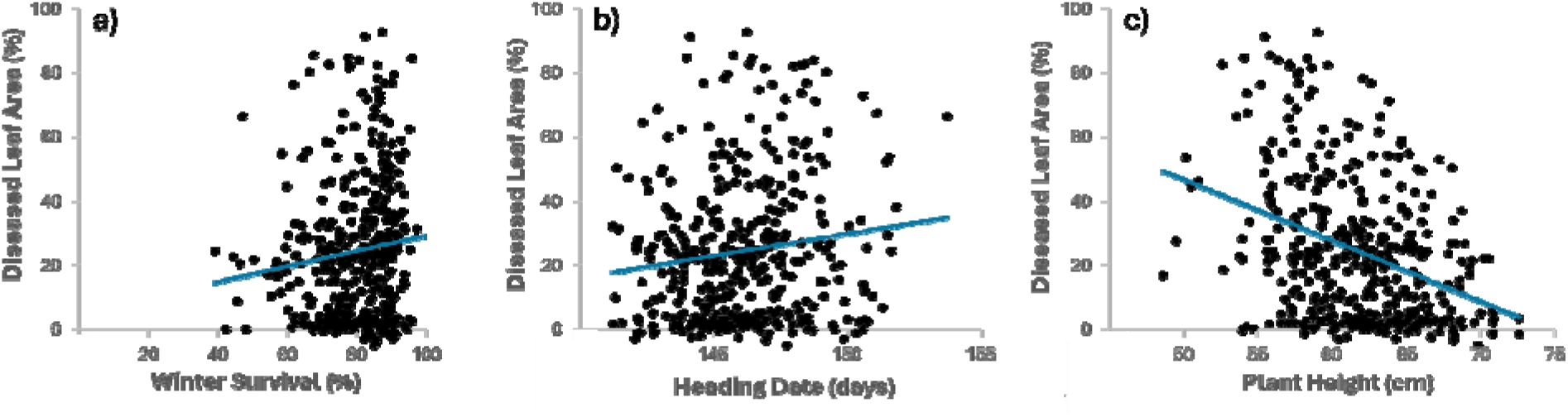
Spearman’s correlation between resistance to scald (diseased leaf area), and a) winter survival, b) heading date and c) plant height, in the 377 line multiparent population across four environments in 2022 and 2023.

**Figure 3.**
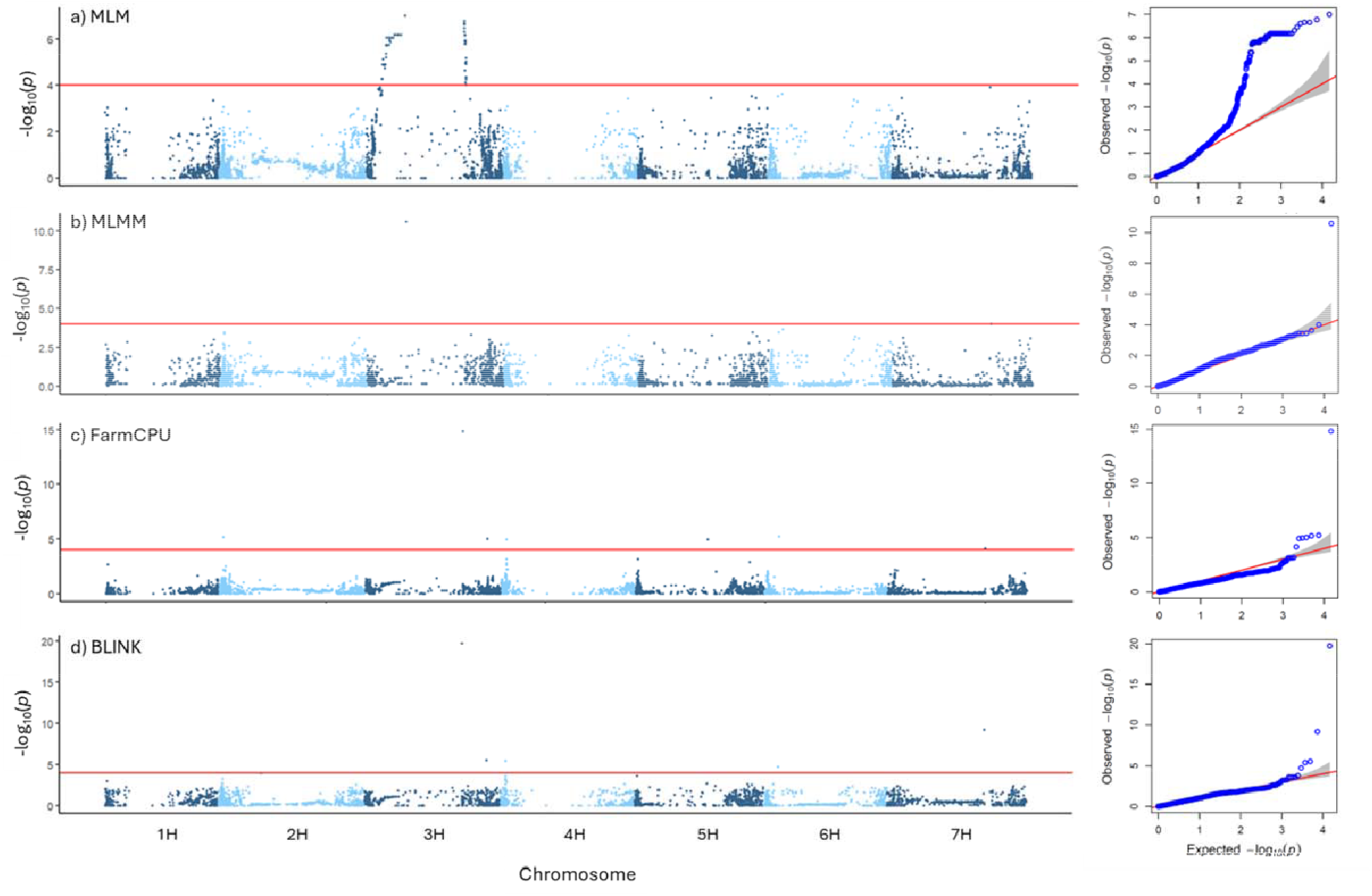
Manhattan plots of the GWAS models including a) MLM, b) MLMM, c) FarmCPU, and d) BLINK, with corresponding QQ plots (right) in GAPIT version 3 for resistance to scald in the multiparent population with using DLA BLUPs for resistance to scald. DLA BLUPs for resistance to scald were calculated using winter survival, heading date and plant height as covariates in the BLUP model in four environments across 2022 and 2023. The red horizontal line in the Manhattan plots indicate the QQ plot - determined threshold of -log_10_(*p*) of 4.0.

#### 2.3.3 GWAS for resistance to scald, winter survival, heading date and plant height

The Box-Cox transformed DLA BLUPS (without WS, HD, and HT as covariates), as well as Box-Cox transformed WS, HD, and HT BLUPs were used as independent trait variables in both MLM and FarmCPU analysis using GAPIT version 3 in RStudio, with the 15,463 SNPs. The significant thresholds were based on visual inspection of the QQ plots of FarmCPU and set to -log_10_(*p*) = 4.0. As in the first GWAS analysis listed above, the models were run again using the ‘N.sig = n’ prompt to determine the correct PVE in GAPIT version 3.

The LD along chromosome 2H was compared to the significant SNP in the marker-trait association for resistance to scald, to determine if there was co-localization and/or LD between WS, HD, and HT to resistance to scald. The Manhattan plot was created in ggplot2 to map the LD and SNP marker trait associations from the MLM GWAS model, and included the clustered SNP regions of the WS, HD and HT marker trait associations on the short end of chromosome 2H. Regions around the DLA, WS, HD and HT SNP associations were based on the SNP associations in the MLM model that were at or above -log_10_(*p*) = 4.0, as an indicator of the confidence interval for the trait. All Manhattan plots were created using ggplot2 in RStudio.

## 3 RESULTS

### 3.1 Segregation for disease and agronomic traits in the multiparent population

Distributions of disease and agronomic traits showed transgressive segregation for most traits from the parental lines (Supplemental Table S1). The distribution of scald DLA was skewed towards the y-axis, with ‘Lightning’ as resistant, ‘SY Tepee’ as moderately resistant, and ‘WintMalt’, ‘KWS Scala’ and ‘Flavia’ as susceptible to scald (Supplemental Fig1). An analysis of variance indicated significant variation for DLA for genotypes, environment and rows within environment. The means and ranges of disease severity for the 377 lines grown across the locations varied with 50.8% (0-99%) in Helfer Farm, 2022, 19.9% (0-99%) in Snyder Farm, 2022, 6.0% (0-90%) in Ketola Farm, 2023, and 19.7% (0-95%) in McGowan Farm, 2023.

Across all three agronomic traits, significant variation was identified for genotype, rows within environment, and columns within environment, but not between environments. Across the four environments, WS was skewed towards 100% survival, with ‘SY Tepee’, ‘WintMalt’ and ‘KWS Scala’ showing good winter survival (>85%). ‘Lightning’ and ‘Flavia’ showed moderate winter survival (∼75-80%), while a check cultivar, ‘Endeavor’, showed poor winter survival. The average winter survival within the population ranged from 75% (Helfer Farm, 2022) to 87% (Ketola Farm, 2023) with an average across the four environments of 79.9%. Differences in heading date between the parents ranged from ∼143 days (Flavia) to ∼151 days (‘WintMalt’), with ‘Lightning’, ‘SY Tepee’ and ‘KWS Scala’ showing mid-range heading dates between 145 days to 147 days. The heading date (Julian) ranged from an average of 142 days in McGowan Farm, 2023 to 150 days to heading in Helfer Farm, 2022. Plant height ranged from an average of 52 cm to 67 cm in Ketola Farm, 2023 (Supplemental Table S1 & S2). Heritability based on an entry-means basis for DLA, WS, HD and HT were 0.76, 0.69, 0.83 and 0.57 respectively.

### 3.2 Scald is correlated with agronomic traits

Correlations between scald DLA and agronomic traits were ascertained to determine the effect of winter survival, heading date and plant height on scald using the 377 lines in the multiparent population (Supplemental Table S2). A highly significant negative correlation was identified between DLA and plant height (*P* < 0.0001; *R* = -0.29), indicating that taller plants had less disease in the upper canopy. A significant correlation was also identified between heading date and scald DLA (*P* < 0.0107; *R*^*2*^ = 0.13), indicating that a later heading date had more DLA. The correlation between winter survival and DLA was not significant across environments (*P* = 0.1065). Notably, there was a significant interaction between winter survival and heading date (*P*=0.0229; *R*^*2*^ = -0.1171), and a highly significant interaction between WS and plant height (*P*<0.0001; *R*^*2*^ = 0.38), and no significant correlation between heading date and plant height.

### 3.3 Genome wide association mapping for resistance to scald

#### 3.3.1 GWAS for resistance to scald using multiple GWAS models

Population structure determined via principal component analysis in GAPIT version 3 (Wang & Zhang, 2021) indicated that the multiparent population consisted of four groupings with the use of only the first and second PCAs, with the top 3 PCA’s accounting for 34% of the genetic variation, indicative of the relatedness between RILs and DHs (Supplemental Figure S2). The QQ plot of the MLM GWAS model showed marked deviation of the observed from expected *p*-values for the MLM model that accounts for both population structure and kinship (Figure 3). The MLMM, FarmCPU and Blink models had QQ plots that were largely in line with the observed and expected *p*-values. The FarmCPU model with PCA = 3 as a covariate resulted in a QQ plot with the observed -log_10_(*p*) values skewing towards the x-axis in comparison to the FarmCPU model with no PCA covariate and was not further pursued (Supplemental Figure S3). The Bonferroni threshold of -log_10_(*p*) = 5.5 was initially used in GAPIT version 3 as the default to determine significance. Analysis of the QQ plots, and particularly the QQ plots from the FarmCPU analysis, showed that a threshold of -log_10_(*p*) = 4.0 was more appropriate and reduced Type II errors, where the observed veered from the expected regression line. Re-analysis of the GWAS models for MLM, MLMM, FarmCPU and Blink threshold set to -log_10_(*p*) of 4.0 was used to estimate the appropriate phenotypic variation explained (Table 1) and showed distinct differences between models (Supplemental Table S3 – S7).

**Table 1.** SNP-trait associations for resistance to scald using DLA BLUPs (using winter survival, heading date and plant height as covariates in the BLUPs), 374 lines and 14,789 SNPs in the MLM, MLMM, FarmCPU and BLINK GWAS models using a threshold of -log_10_(*p*) > 4.0.

| SNP Marker Name | Chromosome | Position (bp)<sup>a</sup> | *p*-value | MAF<sup>b</sup> | Allele | Effect<sup>c</sup> | Phenotype Variance Explained (%) |
| --- | --- | --- | --- | --- | --- | --- | --- |
| **MLM** |  |  |  |  |  |  |  |
| JHI-Hv50k-2016-168548 | 3 | 174,868,558 | $1.00 \times 10^{-07}$ | 0.465 | c/t | -0.0388 | 14.6 |
| SCRI_RS_221644 | 3 | 442,185,927 | $2.21 \times 10^{-07}$ | 0.471 | a/g | -0.0370 | 0.0 |
| JHI-Hv50k-2016-183351 | 3 | 442,550,473 | $4.95 \times 10^{-07}$ | 0.473 | c/t | -0.0358 | 7.1 |
| JHI-Hv50k-2016-183832 | 3 | 446,494,722 | $6.75 \times 10^{-07}$ | 0.472 | t/a | -0.0354 | 7.4 |
| JHI-Hv50k-2016-183828 | 3 | 446,498,335 | $7.21 \times 10^{-07}$ | 0.476 | t/c | -0.0354 | 7.0 |
| **MLMM** |  |  |  |  |  |  |  |
| JHI-Hv50k-2016-168548 | 3 | 174,868,558 | $2.57 \times 10^{-11}$ | 0.465 | c/t | -0.0400 | 22.0 |
| JHI-Hv50k-2016-514374 | 7 | 443,811,200 | $9.90 \times 10^{-05}$ | 0.326 | g/k/n | -0.0434 | 24.1 |
| **FarmCPU** |  |  |  |  |  |  |  |
| SCRI_RS_221644 | 3 | 442,185,927 | $1.55 \times 10^{-15}$ | 0.471 | a/g | -0.0320 | 22.4 |
| JHI-Hv50k-2016-70026 | 2 | 18,144,891 | $6.76 \times 10^{-06}$ | 0.425 | g/a | -0.0120 | 0.0 |
| JHI-Hv50k-2016-202358 | 3 | 553,288,836 | $9.74 \times 10^{-06}$ | 0.300 | t/c | 0.0155 | 15.6 |
| JHI-Hv50k-2016-232214 | 4 | 22,416,684 | $1.20 \times 10^{-05}$ | 0.240 | a/t | 0.0129 | 2.2 |
| JHI-Hv50k-2016-514374 | 7 | 443,811,200 | $7.11 \times 10^{-05}$ | 0.326 | g/t | -0.0308 | 0.0 |
| **BLINK** |  |  |  |  |  |  |  |
| SCRI_RS_221644 | 3 | 442,185,927 | $1.85 \times 10^{-20}$ | 0.471 | a/g | -0.0351 | 7.3 |
| JHI-Hv50k-2016-514374 | 7 | 443,811,200 | $6.54 \times 10^{-10}$ | 0.326 | g/t | -0.0481 | 6.8 |
| JHI-Hv50k-2016-202358 | 3 | 553,288,836 | $2.97 \times 10^{-06}$ | 0.300 | t/c | 0.0163 | 2.7 |
| JHI-Hv50k-2016-232214 | 4 | 22,416,684 | $4.08 \times 10^{-06}$ | 0.240 | a/t | 0.0129 | 3.3 |

<sup>a</sup> SNP position based on Morex version 3 (Mascher et al., 2021)
<sup>b</sup> MAF, minor allele frequency
<sup>c</sup> Effect is based on Box Cox transformed data and as a result is a) not directly transferrable to unit measurements of DLA and b) the negative Box Cox transformation of DLA results in the reverse effect direction of the DLA effect.

##### MLM

The mixed linear model, with population structure and kinship included, resulted in 75 SNPs associated with resistance to scald (Figure 3) that localized in two SNP clusters on chromosome 3H spanning a region from 89 Mbp to 449 Mbp. Two peaks across this region were identified as the significant associations, with the most significant SNP association located at 174,868,558 bp, with a - log_10_(*p*) = 7.0 (MAF = 0.44; PVE = 14.6%. The second most significant SNP was the SCRI_RS_221644 SNP located on the second peak at 442,185,927 bp with a -log_10_(*p*) = 6.8 (MAF = 0.47; PVE = 0.0%). Of the top five SNP associations, four were identified near the second peak.

##### MLMM

Using MLMM, two SNPs were found to be associated with resistance to scald (Figure 3). The single peak identified was located on chromosome 3H, at 174,868,558 bp, with a-log_10_(*p*) = 10.6 (MAF = 0.47; PVE = 22.0%). The second association was identified on chromosome 7H at 443,811,200, with a - logP = 4.0 (MAF = 0.33; PVE = 24.1%).

##### FarmCPU

The FarmCPU model identified five SNPs associated with resistance to scald (Table 1). The most significant SNP was located on chromosome 3H at 442,185,927 bp with a -log_10_(*p*) = 14.8 (MAF = 0.47; PVE = 22.4%). A second SNP on chromosome 3H was located at 553,228,836 with a -log_10_(*p*) = 5.0 (MAF = 0.30; PVE = 15.6%). The SNP association on chromosome 2H was at 18,144,981 bp with a - log_10_(*p*) = 5.2 (MAF = 0.43; PVE = 0.0).Additional SNP associations were identified on chromosomes 4H (MAF = 0.240; PVE = 2.2%) and 7H (MAF = 0.33; PVE = 0.0%). Two additional SNP marker trait associations were identified however both had low MAF and are considered aberrations, including the association on chromosome 6H at 65,635,245 bp (MAF = 0.003; PVE = 0.0%) and on chromosome 5H at 329,813,939 bp (MAF = 0.02; PVE = 5.7%). FarmCPU with PCA=3 resulted in a QQ plot with the observed -log_10_(*p*) values below the expected -log_10_(*p*) values and skewed towards the x-axis and was not considered further (Supplemental Figure S3).

##### BLINK

BLINK identified four SNPs associated with resistance to scald. The most significant SNP was located on chromosome 3H: 442,185,927 bp, with a -log_10_(*p*) = 19.7 (MAF = 0.47; PVE = 7.3%). The next most significant SNP was located on chromosome 7H: 443,811,200 bp with a -log_10_(*p*) = 9.2 (MAF = 0.33; PVE = 6.8%). Other SNP marker trait association included SNPs located on chromosome 3H at 553,288,836 bp with a-log_10_(*p*) = 5.5 (MAF = 0.30; PVE = 2.7%), and 4H at 22,416,684 bp with a -log_10_(*p*) = 5.4 (MAF = 0.24; PVE = 3.3%). As with the FarmCPU model, a SNP association was identified on chromosome 6H: 65,635,245 bp with a -log_10_(*p*) of 4.7 (MAF = 0.003; PAV = 62.3%) and is considered an aberration.

A discrepancy between the GWAS models was identified that was important for interpreting how to proceed with the analysis. The difference in SNP marker trait associations between the MLM, MLMM, FarmCPU and BLINK spans both SNP location and effect size. Notably, there appeared to be two large peaks on chromosome 3H (MLM) that were either an association on the first peak (MLMM) or on the second peak (FarmCPU; BLINK). In addition, PVE for the large peaks on chromosome 3H (442 Mbp) varied between MLM (0.0%), FarmCPU (22..4%), and BLINK (7.3%); the corresponding SNP in MLMM at 174Mbp (22.0%), indicating the presence of a major gene such as the *Rrs1* co-localizes to that region.

Using TASSEL 5.0 (Bradbury et al., 2007), the LD along chromosome 3H was compared to both peaks separately (Figure 4). Using the SNP at 174,868,558 bp, the LD analysis showed that SNPs across both peaks were in LD with the peak at 174 Mbp, indicating a large linkage block that spanned the centromere. The LD analysis comparing the SNPs at the second peak at 442,185,927 bp also indicated that SNPs across both peaks were in LD with the second peak. In both scenarios of GWAS using MLM, where both peaks presented as being associated with resistance, the LD patterns were tightly linked and in coordination with the *p*-values of the surrounding SNP association landscape. As this population is derived from RILs and/or DHs, it appears that the two large peaks on chromosome 3H comprise one large linkage block that spans the centromere. The SNP at 442 Mbp is also the most significant SNP identified via GWAS MLM, and in FarmCPU and BLINK, with FarmCPU indicating the most variation explained by this SNP.

**Figure 4.**
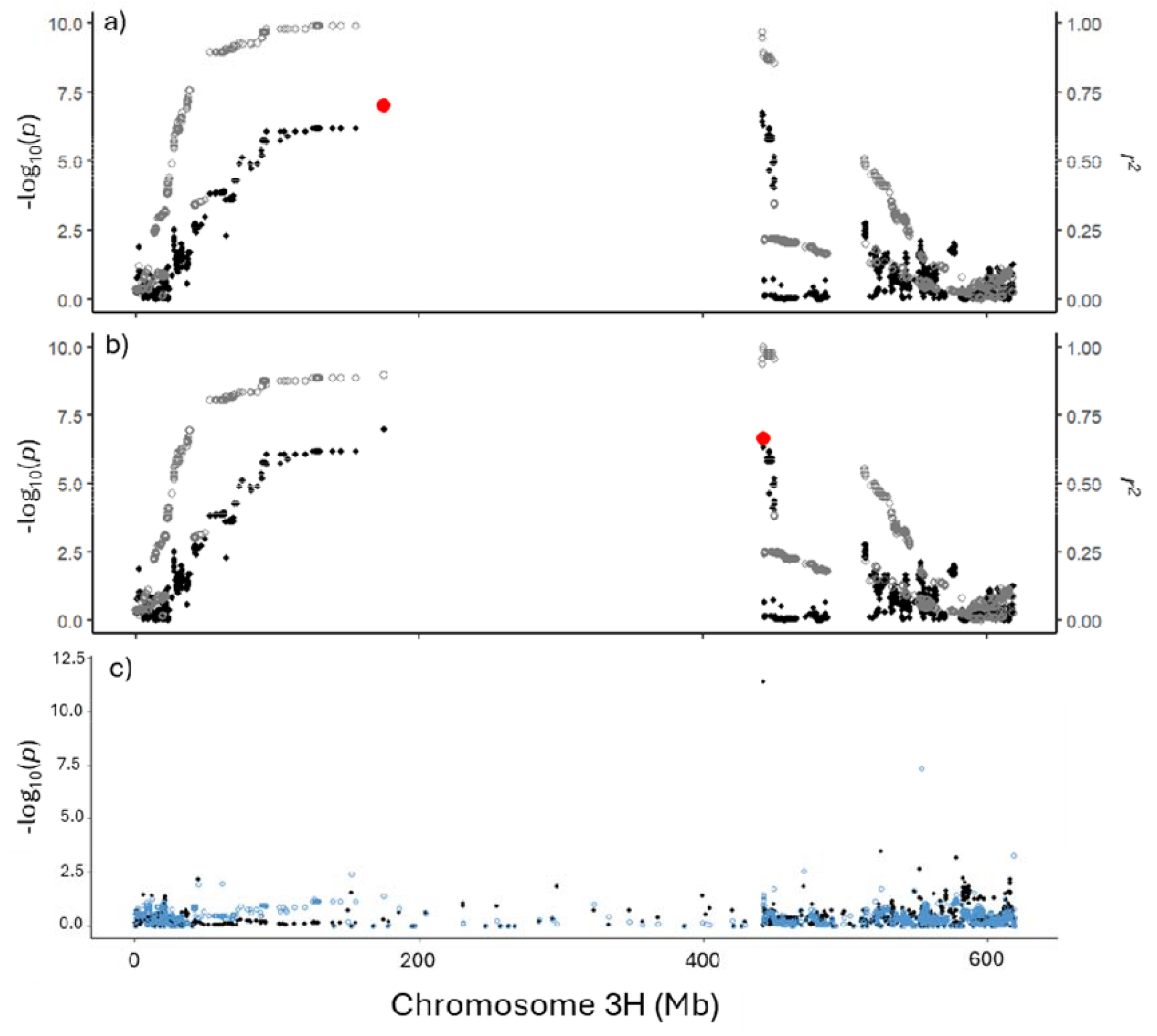
Dissection of the SNP-trait association cluster on chromosome 3H for resistance to scald DLA, where a) characterizes linkage disequilibrium (LD) in relation to the MLM SNP trait association at 174 Mbp, b) characterizes LD in relation to the MLM SNP-trait association at 442 Mbp. SNP associations are labeled as black dots. Linkage disequilibrium indicated as *r*^*2*^ is shown in grey dots in a) and b). The most significant SNP-trait association at 174 Mbp and 442 Mbp chromosome 3H is indicated as a red dot. The FarmCPU SNP associations (c) across chromosome 3H characterizes associations across chromosome 3H in black dots. Associations using the HVS3 (*Rrs1*) SCAR marker as a covariate (c) in blue dots indicates the presence of *Rrs1* at 442 Mbp, and reveals an association at 553Mbp.

#### 3.3.2 *Rrs1* is segregating in a large linkage block on chromosome 3H

The cluster of SNPs with high marker-trait associations on chromosome 3H co-localized near the *Rrs1* locus. To determine if the resistance in this region is due to the major resistance gene complex, *Rrs1*, the population was scored for the HVS3 SCAR marker, located downstream of the *Rrs1* gene (Genger et al., 2003). ‘Lightning’ harbored the resistance allele at 250 bp, while ‘Flavia’, ‘KWS Scala’, ‘SY Tepee’ and ‘WintMalt’ carried the susceptible allele at ∼500 bp. (Supplemental Table S8). The HVS3 marker was used as a covariate in a FarmCPU GWAS analysis in GAPIT version 3 with 354 lines (Supplemental Tables S9 & S10). The Manhattan plot along chromosome 3H (Figure 4c) indicates the presence of the SNP at 442 Mbp without the HVS3 SCAR marker as a covariate, and the absence of the same SNP association when the HVS3 SCAR marker is used as a covariate, indicating the resistance identified via GWAS model is conferred via *Rrs1*. In addition, the use of the HVS3 SCAR marker as a covariate in FarmCPU GWAS model revealed the SNP association at 553 Mbp in this subset of lines, validating the second SNP on chromosome 3H.

Based on the Morex version 3 genome, the SNP at 442,185,927 bp was located within a receptor-like kinase (HORVU.MOREX.r3.HG0281210.1), that is one of several kinase and receptor-like kinase genes clustered in the region. The SNP located on chromosome 3H at 553,288,836 bp co-localized with the *Rrs4* gene (Patil et al., 2003), located at 576.6 Mbp. The SNP resides in a Protein 1Q-Domain 1 gene (HORVU.MOREX.r3.3HG0303590.1). The *Rrs17* resistance gene is located at 10.4 Mbp (Wagner et al., 2008), upstream of the SNP identified on chromosome 2H located at 18.1 Mbp, that is located within a leucine-rich repeat receptor-like protein kinase (HORVU.MOREX.r3.2HG0104570). The identified SNP on chromosome 4H at 22,416,684 bp is located within HORVU.MOREX.r3.4HG0338290, a glucan endo-1 3-beta glucosidase gene. The SNP association on chromosome 7H at 443,811,200 is located within HORVU.MOREX.r3.7HG0707540, a glycosyltransferase gene.

Four GWAS models were used to validate SNP associations across the genome for resistance to scald. All four models have different approaches for estimating associations and are available to avoid overfitting analysis. The most significant SNP association(s) in this study resided on chromosome 3H at either 174.8 Mbp or 422.2 Mbp. FarmCPU and Blink associations at these peaks had the lowest *p*-values (i.e., the most significant) across the models. Importantly, the PVE differed between the models for these two SNPs. The PVE at 174.8 Mbp ranged from 14.6% to 22.0% for MLM and MLMM, respectively. The PVE at 442.2 Mbp was 0.0%, 22.4% and 7.3% for the MLM, FarmCPU and BLINK models, respectively. The FarmCPU model PVE of 22.4% is indicative of major gene resistance such as *Rrs1*. The SNP association on chromosome 6H, explained 62.3% of the phenotypic variation in BLINK despite the very low MAF (0.004) and given the variation in the few SNPs at this locus, appears to be a Type I error. The FarmCPU model constrains the PVE of this SNP at 0.0%. Due to the PVE allocated for the *Rrs1* region, the additional SNPs across the genome such as that on chromosome 3H at 553 Mbp, the lesser PVE of the aberrant SNP on chromosome 6H, and for simplicity, the FarmCPU was the model that was used in the further analysis, being considered the best fit for this population. The MLM model was also used for reference.

#### 3.3.3 GWAS for resistance to scald compared to agronomic traits via FarmCPU

Association analysis across disease and agronomic traits for the 374 lines was used to identify SNP associations for resistance to scald, winter survival, heading date and plant height to determine if there was any colocalization of marker associations that explained the phenotypic traits (Table 2; Supplemental Tables S11 - 18). The BLUPs were estimated as mentioned above and included the scald DLA BLUP estimate (without using the WS, HD, or HT as covariates). The QQ plots for the FarmCPU analysis (Figure 5) indicated an adherence of the observed to expected -log_10_(*p*) values except for several SNPs at the tail of the slope, as expected. The separation of the observed to expected levels at the thresholds for each of the traits was analyzed separately, and based on the Q plots, a -log_10_(*p*) = 4.0 was used as the threshold for each trait (Figure 5).

**Table 2.** SNP-trait associations for diseased leaf area for scald, winter survival, heading date and plant height BLUPs, for the 374 lines and 14,789 SNPs in the FarmCPU GWAS model using a threshold of -log_10_(*p*) > 4.0. using the FarmCPU model.

| SNP | Chromosome | Position (bp)<sup>a</sup> | *p*-value | MAF<sup>b</sup> | Allele | Effect<sup>c</sup> | Phenotype Variance Explained(%) |
| --- | --- | --- | --- | --- | --- | --- | --- |
| **Scald DLA** |  |  |  |  |  |  |  |
| SCRI_RS_221644 | 3 | 442,185,927 | $1.48 \times 10^{-15}$ | 0.471 | a/g | -0.0328 | 29.4 |
| JHI-Hv50k-2016-70026 | 2 | 18,144,891 | $1.14 \times 10^{-06}$ | 0.425 | g/a | -0.0145 | 1.8 |
| JHI-Hv50k-2016-232214 | 4 | 22,416,684 | $7.98 \times 10^{-05}$ | 0.240 | a/t | 0.0103 | 4.8 |
| **Winter survival** |  |  |  |  |  |  |  |
| JHI-Hv50k-2016-61826 | 2 | 4,217,244 | $1.41 \times 10^{-40}$ | 0.432 | a/g | -39533.6 | 0.8 |
| JHI-Hv50k-2016-9518 | 1 | 9,315,052 | $9.45 \times 10^{-09}$ | 0.161 | g/a | -17465.1 | 5.8 |
| JHI-Hv50k-2016-293098 | 5 | 84,325,845 | $1.90 \times 10^{-06}$ | 0.318 | c/t | 22356.3 | 0.3 |
| SCRI_RS_103215 | 3 | 563,918,935 | $3.61 \times 10^{-05}$ | 0.353 | a/g | -9924.2 | 1.8 |
| **Heading date** |  |  |  |  |  |  |  |
| JHI-Hv50k-2016-267990 | 4 | 592,022,994 | $2.41 \times 10^{-17}$ | 0.395 | c/t | -0.4673 | 0.1 |
| JHI-Hv50k-2016-73663 | 2 | 26,107,957 | $1.70 \times 10^{-13}$ | 0.389 | g/t | 0.7865 | 14.0 |
| SCRI_RS_231735 | 1 | 417,457,946 | $7.00 \times 10^{-12}$ | 0.340 | c/a | 0.3784 | 4.7 |
| JHI-Hv50k-2016-456393 | 7 | 31,220,855 | $4.88 \times 10^{-09}$ | 0.150 | g/t | -0.4118 | 0.0 |
| JHI-Hv50k-2016-9518 | 1 | 9,315,052 | $9.34 \times 10^{-09}$ | 0.161 | a/g | -0.3727 | 0.0 |
| JHI-Hv50k-2016-332486 | 5 | 522,212,970 | $6.81 \times 10^{-07}$ | 0.361 | t/a | -0.3034 | 0.0 |
| JHI-Hv50k-2016-331267 | 5 | 519,940,617 | $1.34 \times 10^{-06}$ | 0.200 | a/g | -0.3209 | 0.0 |
| JHI-Hv50k-2016-320694 | 5 | 496,625,681 | $1.59 \times 10^{-05}$ | 0.367 | a/g | -0.2620 | 2.7 |
| JHI-Hv50k-2016-88794 | 2 | 117,994,190 | $5.03 \times 10^{-05}$ | 0.216 | a/c | -0.5310 | 1.0 |
| **Plant height** |  |  |  |  |  |  |  |
| JHI-Hv50k-2016-64130 | 2 | 10,119,392 | $4.81 \times 10^{-14}$ | 0.288 | g/r | 0.0661 | 10.7 |
| JHI-Hv50k-2016-9209 | 1 | 8,999,409 | $5.21 \times 10^{-08}$ | 0.391 | c/t | 0.0304 | 4.1 |
| JHI-Hv50k-2016-216732 | 3 | 595,758,747 | $8.50 \times 10^{-08}$ | 0.426 | a/c | -0.0441 | 7.4 |
| SCRI_RS_150944 | 3 | 560,042,502 | $1.79 \times 10^{-05}$ | 0.342 | t/c | -0.0201 | 1.4 |
| JHI-Hv50k-2016-281309 | 5 | 9,453,080 | $1.87 \times 10^{-05}$ | 0.465 | c/t | -0.0178 | 5.5 |
| JHI-Hv50k-2016-250342 | 4 | 494,857,832 | $1.94 \times 10^{-05}$ | 0.198 | g/r | 0.0386 | 0.0 |
| JHI-Hv50k-2016-175127 | 3 | 322,820,460 | $5.94 \times 10^{-05}$ | 0.499 | n/y | -0.3041 | 3.8 |

<sup>a</sup> SNP position based on Morex version 3 (Mascher et al., 2021)
<sup>b</sup> MAF, minor allele frequency
<sup>c</sup> Effect is based on Box Cox transformed data and as a result is a) not directly transferrable to unit measurements of traits and b) the negative Box Cox transformation of DLA and HD results in the reverse effect direction of effects.

**Figure 5:**
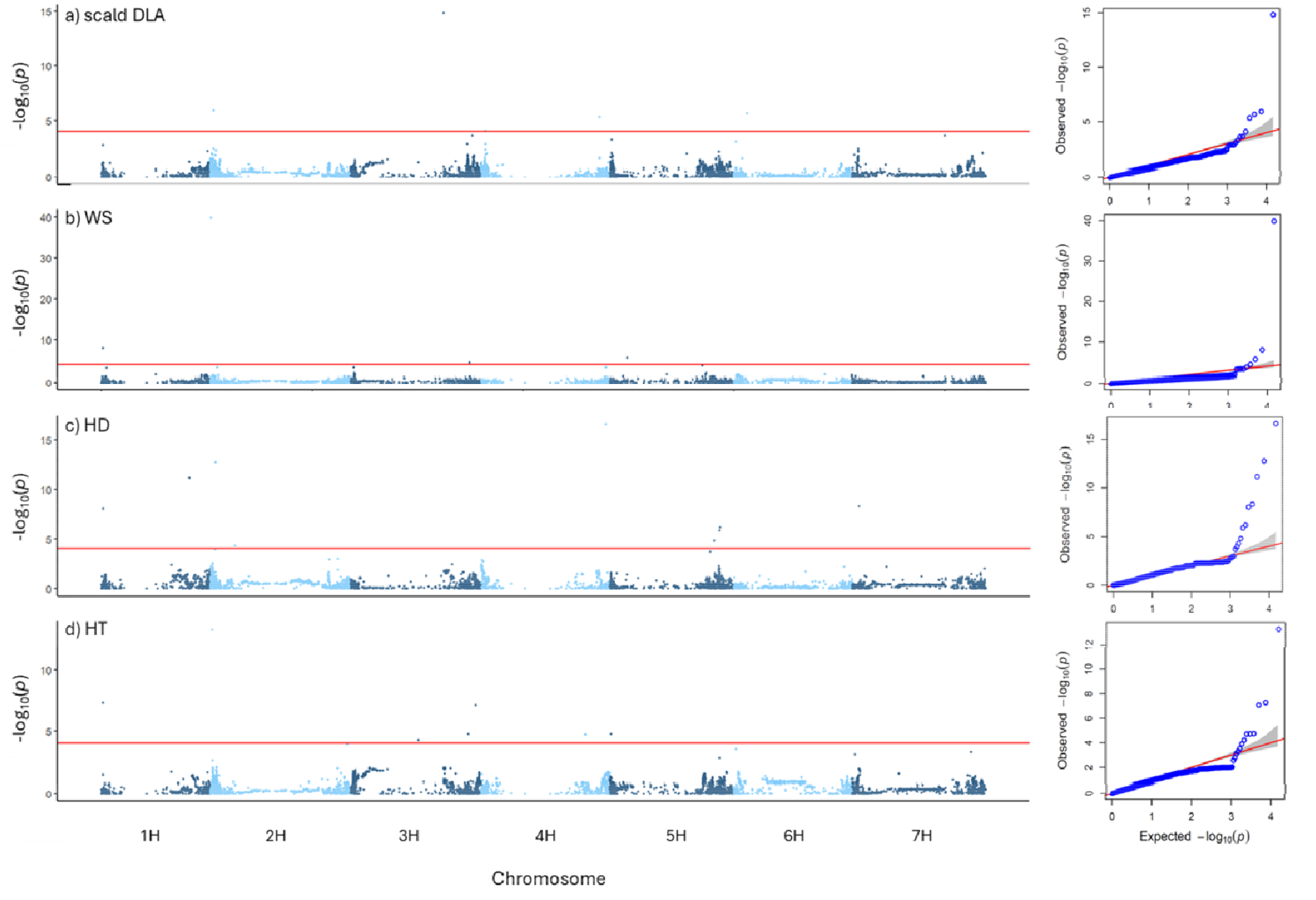
Manhattan plots for a) resistance to scald (DLA), b) winter survival (WS), c) heading date (HD), and d) plant height (HT) using the 374 lines and 14,789 SNPs in the FarmCPU GWAS model, and corresponding QQ plots (right). DLA BLUPs for resistance to scald were calculated without using winter survival, heading date and plant height as covariates in the BLUP model. All traits BLUPs were calculated in four environments across 2022 and 2023. The red horizontal line indicates the QQ plot – determined threshold of -log_10_(*p*) of 4.0.

DLA: In general, the SNP associations identified using the scald DLA BLUPs (without the agronomic traits as covariates) using FarmCPU (Figure 5; Table 2) were similar to the previous identified SNPs (using the agronomic traits as covariates; Table 1). The SNP associations on chromosome 3H included the main association at 442,185,927 bp (PVE=29.4%). The same SNP associations were identified on Chromosome 2H and 4H, with slightly adjusted PVE of 1.8% and 4.8%, respectively. The SNP association aberrations identified in the previous analysis were also identified in this analysis and not considered for further discussion based on low MAF and exaggerated PVE for the SNP on Chromosome 6H.

Winter Survival: A SNP association was identified on Chromosome 2H, at 4,217,244 bp (MAF = 0.432; PVE = 0.8%). Additional SNPs were identified in FarmCPU including SNPs on chromosome 1H at 9,315,052 bp (MAF = 0.161; PVE = 5.8%), chromosome 5H at 84,325,845 bp (MAF = 0.318; PVE = 0.3%), and chromosome 3H at 563,918,935 bp (MAF = 0.353; PVE = 1.8%).

Heading date: Of the nine SNP associations for heading date, the most significant association co-localized at *Ppd-H1* (Turner et al., 2005), on chromosome 2H at 26,107,957 (MAF = 0.389; PVE = 14.0%). The SNP association with the second highest PVE was located on Chromosome 1H at 417,457,946 bp (MAF = 0.340; PVE = 4.7%). Similar to WS, a SNP associations were identified on chromosome 1H at 9,315052 (MAF = 0.161; PVE = 0.0%). The remaining SNP associations contributed <0.1% PVE. Plant Height: Seven SNP associations were identified for HT (Table 1). The most significant SNP, chromosome 2H: 10,011,392 (MAF = 0.288) contributed approximately 10.7% to the phenotypic variation and is localized upstream of *Ppd-H1* and the SNP association peak identified for heading date. A SNP association on chromosome 1H was located at 8,999,409 bp (MAF = 0.391; PVE = 4.1%), just upstream of the associations identified for winter survival and heading date. Two SNPs were associated on chromosome 3H, located at 5995,758,77 bp (MAF = 0.426; PVE = 7.4%) and 560,042,502 bp (MAF = 0.342; PVE = 1.4%). Additional SNP associations were identified on chromosome 3H, 4H and 5H that accounted for PVE from 0.0 to 5.5%,.

The *HvGA20ox2* gene, also known as *sdw1*, is located on chromosome 3H at 563.9 Mbp and, in previous GWAS studies, was suggested to contribute towards reaction of barley to scald (Looseley et al., 2018). In our study, two SNP associations for plant height were located within this region at 560.0 Mbp and 595.7 Mbp. The multiparent population and parental lines were genotyped using the *sdw1.d* and *sdw1.c* markers (Xu et al., 2017), all of which carried the wild type allele at sdw1.

The correlation between resistance to scald and agronomic traits resulted in the co-localization of GWAS SNP associations. The inclusion of agronomic traits in the DLA BLUPs as covariates in the initial analysis (Table 1; Figure 3) appeared to slightly lower the *p*-values for identified SNP associations and resulted in a higher significance threshold in comparison to DLA BLUPs without covariates, with the exception of the SNP association linked to *Rrs1*. The scald DLA association identified at 553.3 Mbp on chromosome 3H, was only identified when the agronomic traits were used as covariates in the scald DLA BLUPs, likely due to the plant height associations found in the region. There were distinct SNP associations on the short arm of chromosome 2H for resistance to scald (chromosome 2H: 18.1 Mbp; Figure 6), winter survival (chromosome 2H: 4.2 Mbp), heading date (chromosome 2H; 26.1 Mbp) and plant height (chromosome 2H: 10.1 Mbp), however there appeared to be little genomic overlap between these peaks, as seen in the Manhattan plots for the GWAS MLM model (Supplemental Fig S4). The linkage disequilibrium between the significant peak for resistance to scald (DLA) and the remaining SNPs along chromosome 2H indicated that this region may harbor LD, and that the scald DLA association should likely be considered a QTL related to architectural avoidance (Figure 6). Alternatively, the SNP association is located within a gene encoding agmatine coumaroyltransferase-2 (ACT-2), known to produce antifungal compounds (Burhenne et al., 2003). The resistance gene, *Rrs17*, is also located at 10.4 Mbp (Wagner et al., 2008).

**Figure 6.**
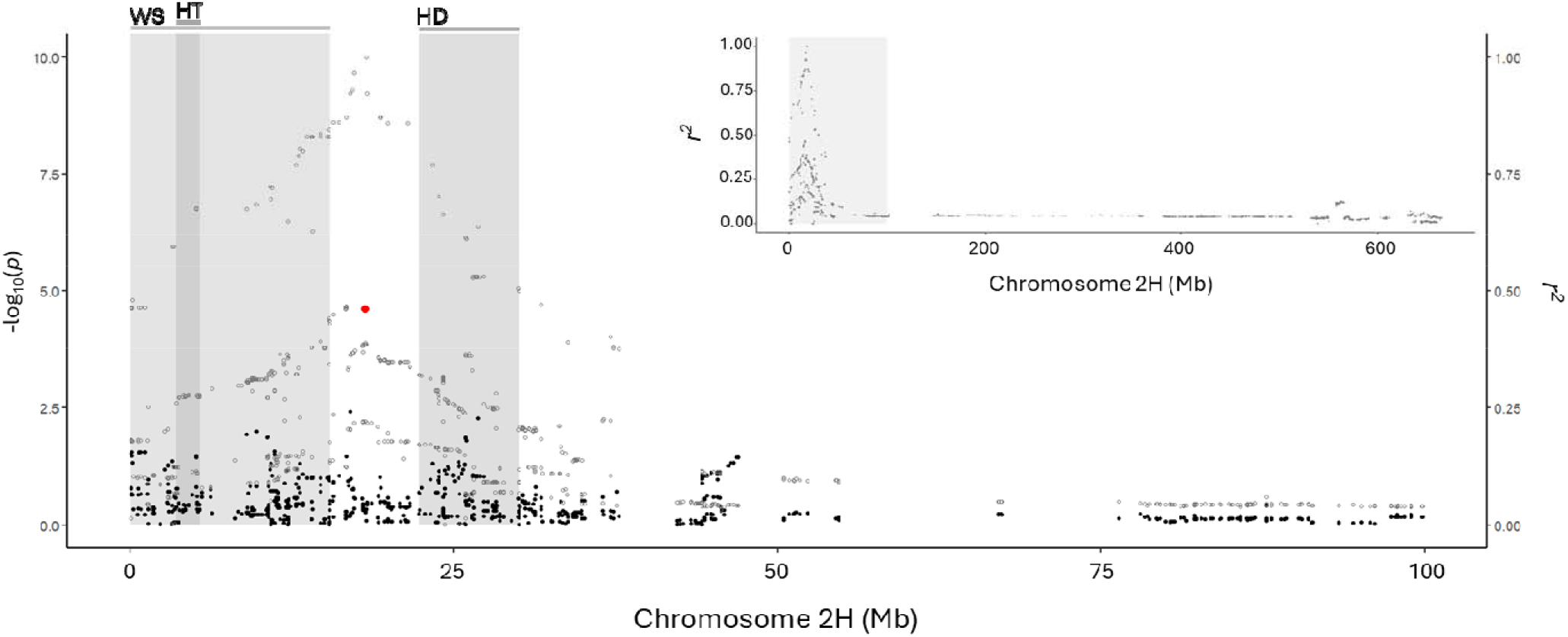
Association and linkage disequilibrium (LD) analysis across chromosome 2H for resistance to scald (DLA; black dots). The Manhattan plot (left y axis) with black dots depicts the SNP associations for resistance to diseased leaf area (DLA) using FarmCPU in GAPIT version 3 with the most significant SNP association identified as a red dot across the first 1 Mbp of chromosome 2H. Grey vertical shaded regions in the main graph represent confidence intervals based on significant MLM associations (Supplemental Figure S) of co-localized SNP-trait association clusters for winter survival (left, grey bar), heading date (right, grey bar) and plant height (left, dark grey bar). The LD estimate scatter plot (right *y-*axis) with grey dots indicates the LD of neighboring SNPs in relation to the SNP associated with resistance to scald (in red). The top right graph insert indicates the LD across chromosome 2H with respect to the most significant associated SNP for DLA. The vertical grey shaded box indicates the 1 Mbp region shown in the main graph.

## 4 DISCUSSION

### 4.1. Breeding for resistance to scald in winter malting barley is important

Malting barley is a commercially important crop due to its value in the brewing and distilling industries. The malting process demands uniformly high quality grain. Winter malting barley, grown in New York for the craft brewing industry, faces unique challenges due to the wet humid weather in this region. Foliar fungal diseases affect both spring and winter malting barley, resulting in yield loss and kernel quality issues. The environmental conditions encountered by winter malting barley grown in New York are more conducive to scald infections than spring barley, even when grown in similar regions (Kolkman et al., 2025a, b).

In this study, ambient inoculum of *R. graminicola* was relied upon for field infections. Winter malting barley is planted in the fall and develops in the early spring season in New York when weather is favorable for development, reaching levels of up to 77% severity in susceptible germplasm (Kolkman et al., 2025a). The distribution of the susceptible parent check, ‘KWS Scala’, across each of the environments in this study validated the consistent natural infection across the experiment, while early season infections in ‘KWS Scala’ provided spore production across the experiment that resulted in consistent disease severity in highly susceptible genotypes.

### 4.2 Resistance in the multiparent population is comprised of both major resistance genes and quantitative disease resistance

The *Rrs1* gene complex is an important source of resistance to scald. We identified a large linkage block on chromosome 3H that spanned the centromere and initially appeared as two separate peaks in the MLM GWAS model. Linkage disequilibrium analysis validated that the large region was in LD with the most significant peak that co-localized near the *Rrs1* gene complex. Additional evidence of the large linkage blocks was seen in the deviation of the observed: expected *p*-values as seen by the heavily skewed slope in the QQ plots for the MLM model. The evidence of a large linkage block at *Rrs1* has been seen in previous GWAS studies, where a large linkage block was identified on chromosome 3H for seedling resistance in a diversity panel of spring barley (Hiddar et al., 2023) which was accompanied by QQ plots that showed a large deviation from the expected, and where there were two peaks in LD that spanned the centromeric region (Kunze et al., 2024).

Fine-mapping *Rrs1* to a 0.8 cM has identified ten candidate genes, however the *Rrs1* allele is in a centromeric region with high linkage disequilibrium and is not present in the reference Morex genome (Looseley et al., 2020). Further association analysis derived three markers near a protein kinase gene, indicating the *Rrs1* may be an allelic a member of the protein kinase gene family (Looseley et al., 2020). Examination of the mode of action of the *Rrs1* gene upon infection reveals that the resistance is incomplete, but that hyphal development is restricted, random in the direction of growth, and exhibits a delay in the collapse of the epidermal layer (Thirugnanasambandam et al., 2011). Additionally, marker trait associations on chromosome 3H have been found to contribute ∼27 to 30% of the phenotypic variation for scald (Kunze et al., 2024; Noe et al., 2025), similar to the 22.4 – 29.4% phenotypic variation explained at *Rrs1* in this study. Importantly, additional SNP marker trait associations were found across the genome, including one that co-localized with *Rrs4* on chromosome 3H, and regions on chromosomes 2H, 4H, and 7H that can add to durable long term resistance.

### 4.3 Agronomic traits can reduce disease infection

*Rhynchosporium graminicola* is a slow growing pathogen, requiring a three-week incubation period between infection and lesion development. It capitalizes on the long early cold days of spring to increase in spore load and infection. Agronomic variables can influence the rate of upward dispersal of spores in the plant canopy. Winter survival is an important trait in crop improvement in northern environments. The positive correlation between reduced plant density from poor winter survival (and resulting reduced plot canopy) and reduced disease is an indicator of a microclimate and putative reduced *R. graminicola* growth and/or spore load that leads to less disease. While not useful for breeding purposes regarding management of scald through reduced canopy density, it is important to include in GWAS models as a covariate for the phenotypic trait to improve SNP association identification.

Additional agronomic traits can help reduce infection. The negative correlation between heading date and scald is indicative of an escape or avoidance mechanism, limiting the ability of *R. graminicola* conidia to reach the upper leaves and spikes. The negative correlation between plant height and diseased leaf area for scald also suggests an avoidance mechanism in plant defense, where elongative growth of plants may outpace pathogen reproductive cycles thus delaying the rain splash dispersal of conidia to upper leaves and glumes. The *sdw1* gene has previously been identified in GWAS for resistance to scald (Loosely et al., 2018). In this study we identified SNP associations for resistance to scald as well as plant height in the region of *sdw1* that may be independent of each other and/or a functional variant of *sdw1* that has not been reported. In addition, a region on the short arm of chromosome 2H contributed to plant height and heading date (via *Ppd-H1*) and to a lesser extent, winter survival. The SNP association in this region may contribute to either plant architectural traits, and/or active resistance, however it provides a target for selection. With a rapidly mutating pathogen, breeding for earlier heading date and taller plants would help provide additional mechanisms for plant defense. The implications for disease management as shorter plants are often selected or desired for reduced lodging should include selection for resistance to scald.

### 4.4 When your population is not that diverse: Selection of GWAS model for adapted multiparent populations

GWAS is a powerful tool for discovery of trait genetic architecture within a population. The identification of SNP marker trait associations can also aid in confirmation of previously identified genetic elements, if the allele is segregating in the population at the appropriate frequency and amplitude to be detected. Understanding the allele effect, or percent variation explained by the SNP/marker trait association can have implications that are important to move forward with more basic genetic studies, or in selection for breeding programs. Most GWAS studies utilize diverse genetic populations for the pursuit of trait association analysis, relying on the thousands of meiotic recombination events between the diverse lines to determine the precise genetic location of marker trait associations. The availability of high-density genotyping platforms for breeding programs offers the ability to use GWAS for crop improvement (Spindel et al., 2013). Breeding programs often cycle many semi-related genetic materials through preliminary breeding trials to select a set number of lines to move forward for selection. While early generation selection of specifically targeted and agronomically important high heritability traits may limit the germplasm pool, the preliminary breeding trials are often replicated across locations. Harnessing the power of GWAS in large breeding populations can be very useful for understanding the genetic architecture of segregating traits in breeding programs.

The multiparent population was derived from a series of biparental crosses and presented a challenge in how to separate population structure from effect. In this study, the SNP associations near *Rrs1* on chromosome 3H varied from 174.9 Mbp accounted for 22.0% in MLMM), whereas the SNP association at 442Mbp accounted for 0.0% (MLM), 22.4% (FarmCPU) and 7.3% (BLINK). The FarmCPU model appeared to be the least likely to overfit or overamplify Type I associations in the structured population. The MLM and MLMM GWAS models include both kinship and population structure whereas FarmCPU includes kinship of associated markers and BLINK includes neither in their respective models. Population structure and kinship are useful tools to reduce the confounding effects and to target unbiased marker trait associations in large diverse populations. FarmCPU detected the largest amount of variation for resistance to scald at the *Rrs1* locus on chromosome 3H, as well as additional QTL using more limited kinship but not population structure.

Breeding for durable resistance to *R. graminicola* should encompass pyramiding a variety of mechanisms to reduce the loss of yield and quality to scald. Major genes, such as *Rrs1*, provide a high level of resistance and can be combined with QTLs and additional major resistance genes, especially those with differing modes of action, to increase the resistance profile and decrease scald. In addition, selection for agronomic traits, such as early flowering time and increased plant height can aid in limiting the vertical spread of disease in the canopy and spike, in order to limit loss of quality and yield to scald.

## Abbreviations

BLINK: Bayesian-information and linkage-disequilibrium iteratively nested keyway
DH: doubled haploid
DLA: diseased leaf area
FarmCPU: fixed and random model circulating probability unification
GAPIT: genome association and prediction integrated tool
GWAS: genome wide association study
HT: height
LD: linkage disequilibrium
LRR-RLK: leucine-rich repeat receptor-like proteins
MAF: minor allele frequency
MAGIC: multiple advanced generation inbred cross
MTA: marker trait association
MLM: mixed linear model
MLMM: multiple loci mixed linear model
PCA: principal components analysis
RIL: recombinant inbred line
SNP: single nucleotide polymorphism
QTL: quantitative trait loci
PEI: pectin esterase inhibitor
QQ: quantile-quantile
Reml: restricted maximum likelihood model
SCAR: sequenced characterized amplified region
PVE: phenotypic variation explained
WS: winter survival

## ACKNOWLEDGEMENTS

The authors thank the field staff at the Cornell University Love Lab and Research Farm, and the small grains field manager and crew, including David Benscher, Jason Schiller, Priscilla Thompson, Jenna Rice and James Tanaka. We also thank Drs. Tyr Wieser-Hanks and Zhiwu Zhang for technical and theoretical help with GAPIT version 3/FarmCPU, respectively. This research was supported by the intramural research program of the U.S. Department of Agriculture, National Institute of Food and Agriculture, Hatch accession #7002745, the USDA Barley Pest Initiative, and the New York State Department of Agriculture and Markets.

## AUTHOR CONTRIBUTIONS

Judith M. Kolkman: Conceptualization; Data curation; Formal analysis; Investigation; Methodology; Visualization; Writing – original draft; Writing – review & editing.

Siim. S. Sepp: Investigation; Writing – review & editing.

Karl Kunze: Conceptualization; Investigation; Writing – review & editing.

Gary C. Bergstrom: Conceptualization; Funding acquisition; Project administration; Supervision; Writing – review & editing.

Mark E. Sorrells: Conceptualization; Funding acquisition; Project administration; Supervision; Writing – review & editing.

## SUPPLEMENTAL MATERIAL

All supplemental figures and tables are available for download upon publication.

## DATA AVAILABILITY

Phenotypic data and GWAS results are available in supplemental tables; genotypic SNP dataset is available at (https://ics.hutton.ac.uk/50k)

## SUPPLEMENTAL FIGURES CONTENT

**Figure S1.**
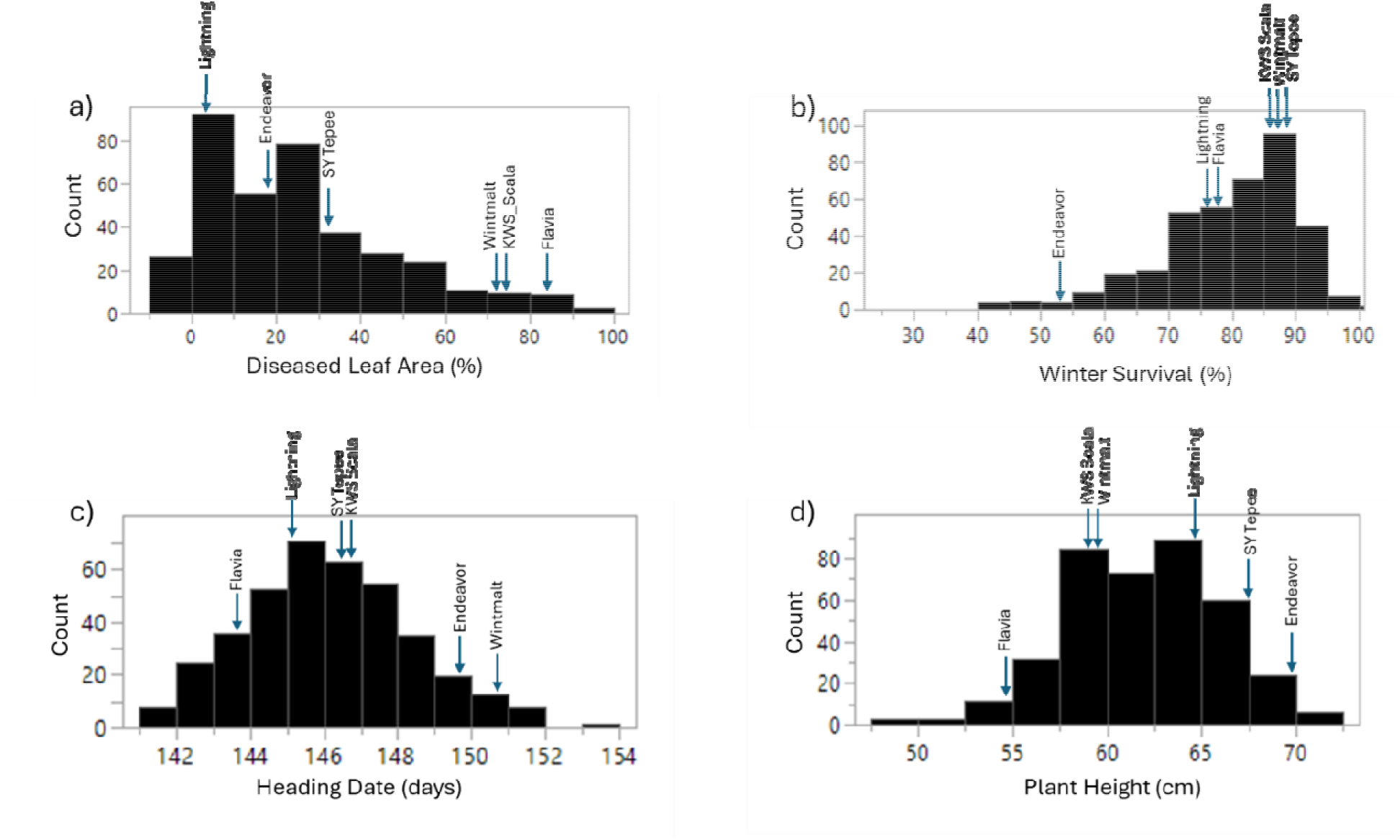
Frequency distribution for a) scald diseased leaf area, b) winter survival, c) heading date (Julian), and d) plant height shows transgressive segregation in the population of 377 lines derived from the parental lines. Values for each parental line are indicated by the blue arrows for each of the disease and agronomic traits. ‘Endeavor’, a check cultivar grown across blocks and experiments is also shown for comparison.

**Figure S2.**
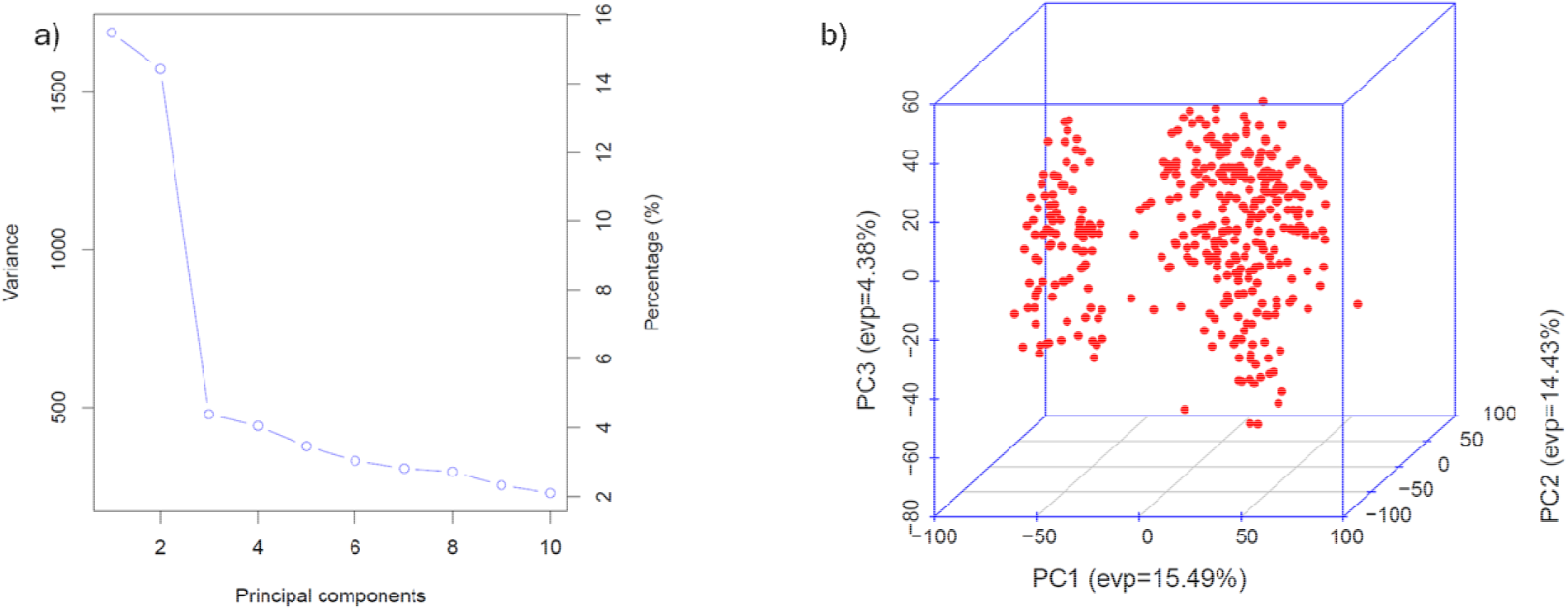
Principal component analysis for the 374 lines of the multiparent population displaying a) PCA eigenvalues and b) genotype clustering for PCA = 3.

**Figure S3.**
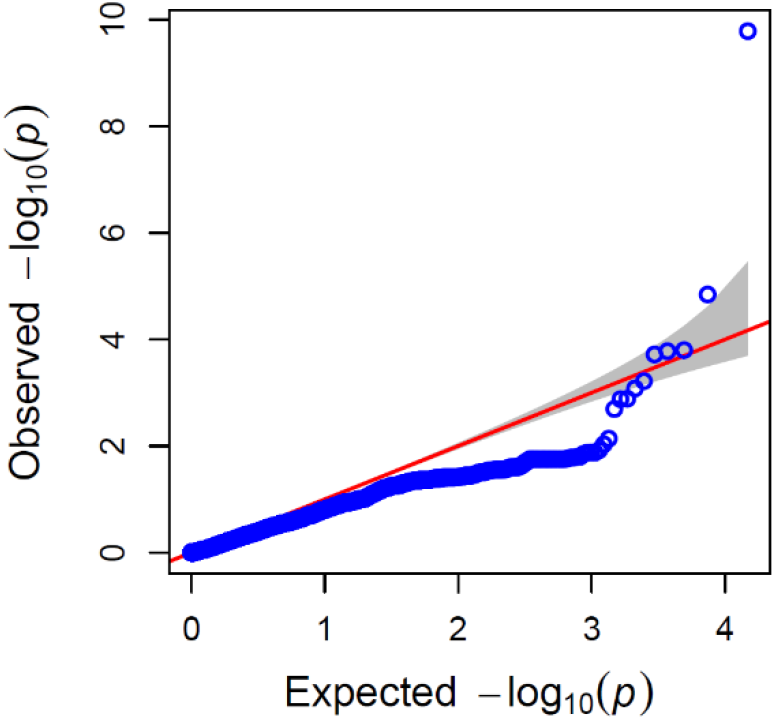
Quantile-quantile plot for FarmCPU GWAS model for resistance to scald in winter malting barley using PCA=3 as a covariate.

**Figure S4:**
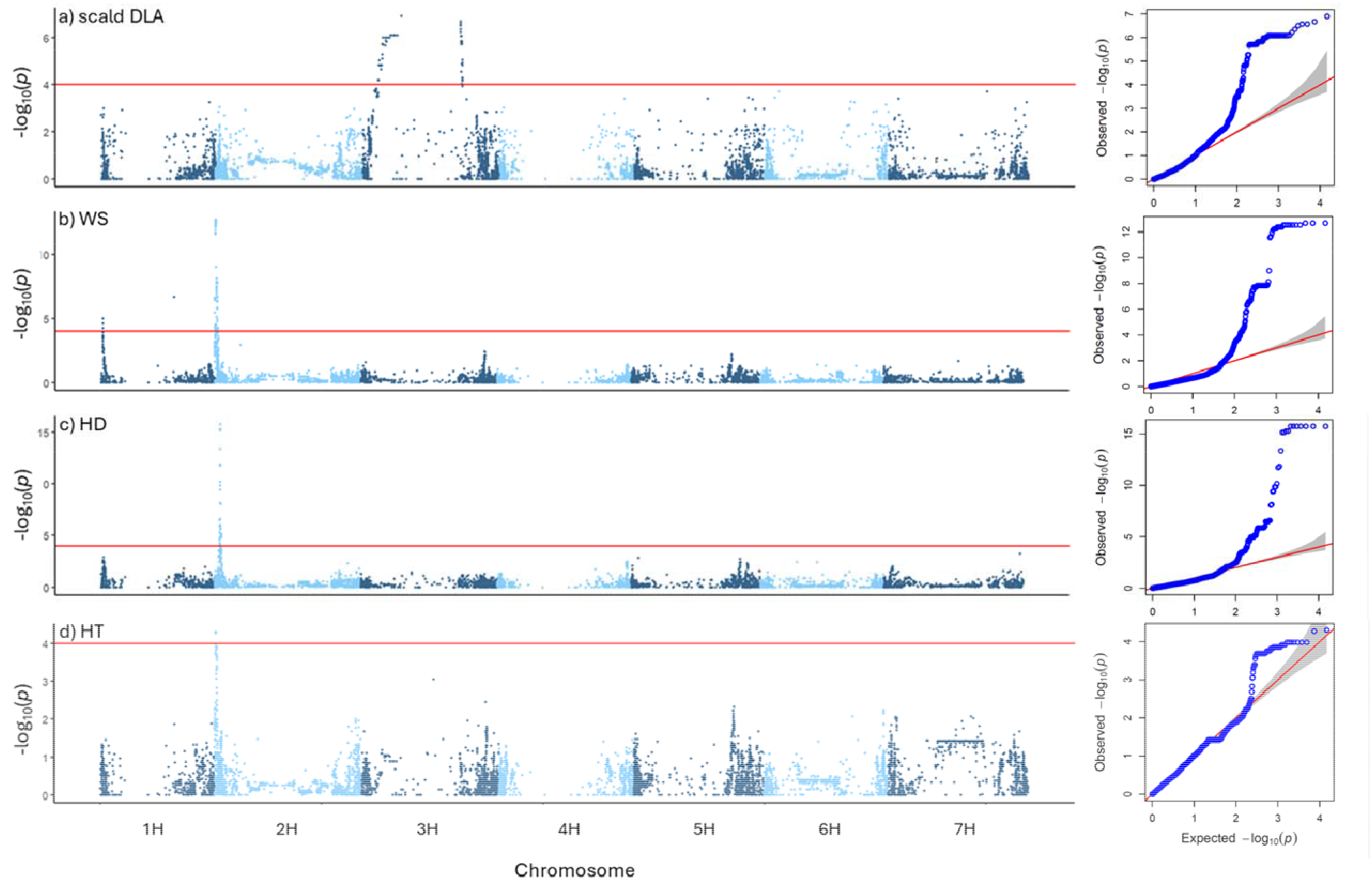
Manhattan plots for a) resistance to scald (DLA), b) winter survival (WS), c) heading date (HD), and d) plant height (HT) using the 374 lines and 14,789 SNPs in the MLM GWAS mode, with corresponding QQ plots (right). DLA BLUPs for resistance to scald were calculated without using WS, HD date and HT as covariates in the BLUP model. All traits BLUPs were calculated in four environments across 2022 and 2023. The red horizontal line indicates the QQ plot – determined threshold of -log_10_(*p*) of 4.0.

## SUPPLEMENTAL TABLES CONTENT

Table S1. Diseased leaf area for scald, winter survival, heading date and plant height in the multiparent population across four environments.

Table S2. Least square means of diseased leaf area for scald, winter survival, heading date and plant height for the multiparent population across four environments.

Table S3. Transformed diseased leaf area BLUPs with and without agronomic covariates, as well as the transformed winter survival, heading date and plant height across four environments.

Table S4. GWAS MLM statistics for DLA BLUPs with agronomic trait covariates

Table S5. GWAS MLMM statistics for DLA BLUPs with agronomic trait covariates

Table S6. GWAS FarmCPU statistics for DLA BLUPs with agronomic trait covariates

Table S7. GWAS BLINK statistics for DLA BLUPs with agronomic trait covariates

Table S8. HVS3 SCAR marker allele sizes linked to *Rrs1* in the multiparent population.

Table S9. GWAS FarmCPU statistics for DLA BLUPs on chromosome 3H with 354 line subset

Table S10. GWAS FarmCPU statistics for DLA BLUPs on chromosome 3H with 354 line subset and HVS3 covariate.

Table S11. GWAS MLM statistics for DLA BLUPs with no agronomic covariates

Table S12. GWAS FarmCPU statistics for DLA BLUPs with no agronomic covariates

Table S13. GWAS MLM statistics for WS BLUPs

Table S14. GWAS FarmCPU statistics for winter survival BLUPs

Table S15. GWAS MLM statistics for heading date BLUPs

Table S16. GWAS FarmCPU statistics for heading date BLUPs

Table S17. GWAS MLM statistics for plant height BLUPs

Table S18. GWAS FarmCPU statistics for plant height BLUPs

